# Inhibition of TYK2 attenuates hyper IL-6- and Oncostatin M-mediated Calcium Signalling in Sensory Neurons

**DOI:** 10.64898/2026.05.15.725418

**Authors:** Thomas Andrew Pritchard, Rohit Gupta, James Higham, Qasim Aziz, David Bulmer

## Abstract

Inflammatory bowel disease (IBD) is characterised by chronic pain, a debilitating symptom for which effective treatments are few and far between. IBD pathogenesis includes the prevalence of a variety of pro-inflammatory cytokines, including the Interleukin-6 (IL-6) family members Il-6 and Oncostatin M (OSM). Previous research has shown disruption of OSM signaling can modulate nociceptor sensitization and activation, however the downstream signalling pathway is unknown. When an *in silico* analysis of murine colonic sensory neuronal populations was undertaken for receptor expression for OSM and other factors necessary for intracellular signaling, we can find diverse expression indicative of functional signaling. We were able to observe that hyper Il-6 (Il-6 bound to the soluble Il-6 receptor) and OSM can elicit activation of a subset of murine sensory neurons by finding an increase in calcium mobilization following superfusion. This could then be attenuated by the pharmacologic inhibition of all janus kinases or interestingly, TYK2 alone. Furthermore, inhibition of transient receptor potential vanilloid 1 or transient receptor potential ankyrin 1 ion channels, which are known to be sensitized by OSM in other sensory neurons also reduced the proportion of OSM-responsive neurons. This further understanding of OSM signaling in sensory neurons creates avenues for more extensive research into the molecular mechanisms occurring as well as the potential to exploit these therapeutically to induce analgesia in a subset of neurons.

## Intro

Inflammatory bowel disease (IBD) is a gastrointestinal disorder primarily comprising of Crohn’s disease (CD) and ulcerative colitis (UC). Whilst these conditions differ in the location and continuation of the inflammatory lesions within the large intestine for which they are known, patients similarly experience a debilitating impact to their daily life (Lee and Lee, 2016; Berre *et al*., 2020). One of the primary correlates in the poor quality of life scores IBD patients present is chronic pain, particularly in the abdomen, and although this is often associated with an episode of inflammation (a “flare”) this can also continue during clinical remission (Zeitz *et al*., 2016; Takahashi *et al*., 2021). Although continued advancement in the development of various immunomodulatory drugs has provided physicians with an arsenal of treatments to combat pathological inflammation in most patients, there remains a dearth of options when it comes to targeting the substantial pain component of the disease (Baille *et al*., 2024; Vieujean *et al*., 2025). Clinicians are wary of prescribing non-steroidal anti-inflammatory drugs due to fears of inducing flares, and although 21% of IBD outpatients in the U.S. use opioids, there is little evidence for effectiveness in chronic abdominal pain as well as potentially exacerbating functional gastrointestinal problems and inducing visceral hyperalgesia (Takeuchi *et al*.,2006; Agostini *et al*., 2010; Kurlander and Drossman, 2014; Bakshi *et al*., 2021; Niccum *et al*.,2021).

The visceral afferent nerve terminals which innervate the large intestine detect noxious chemical and mechanical stimuli, and prolonged exposure to such as in the case of the inflamed colon in IBD, can lead to visceral hypersensitivity even upon the resolution of a flare (Willert *et al*., 2004). Previous studies have shown sensitisation of the ligand-gated ion channel transient receptor potential vanilloid 1 (TRPV1), and the non-selective cation channel transient receptor potential ankyrin 1 (TRPA1) by tumour necrosis factor α amongst other pro-inflammatory mediators abundant in IBD colonic tissue (Lapointe *et al*.,2015; Balemans *et al*., 2017; Barker *et al*., 2022; Higham *et al*., 2024). A cytokine which has been found to be similarly upregulated but lesser explored is Oncostatin M (OSM; West *at el*., 2017).

OSM is a pleotropic cytokine known to induce immune cell differentiation, further cytokine release, and a variety of other functions in a tissue dependant manner (West *et al*., 2018). Through binding with the obligate receptors OSMR -and leukemia inhibitory factor receptor (LIFR)- and GP130, OSM triggers the phosphorylation of the intracellular tyrosine janus kinases (JAK) which exist in a heteromeric or homomeric dimer (Rose-John, 2018). Interestingly, analysis of a scRNA-seq dataset published by Hockley *et al*. (2019) revealed substantial transcript levels of the obligate OSM receptor genes (*Osmr*) and glycoprotein-130 (*Il6st*) in murine colonic-innervating nociceptor populations. Co-expression could be observed between OSMR and GP130 consistent with observations by Langeslag *et al*. (2011) whereby OSM was seen to potentiate capsaicin (TRPV1 agonist) responses in a GP130 dependant manner, highlighting a mechanism by which OSM could activate colonic nociceptors. In a recent publication by Li *et al*. (2026) OSM was found to induce action potential generation and paw mechanical hypersensitivity in rats but found little evidence to suggest any sensitisation to heat responses or co-expression in DRG neurons between TRPV1 and OMSR.

Through the *ex vivo* analysis of hyper IL6 (a fusion protein of Il-6 and the soluble Il-6 receptor) and OSM-induced [Ca^2+^] flux in sensory neuron cell bodies and pharmacologic disruption of intracellular signalling, we seek to determine whether OSM may activate nociceptors in a TYK2 dependant manner, highlighting another potential route for inflammatory-mediated nociception in the context of IBD.

## Methods

### 2.3 Animals

All mice used in experiments consisted of a combination of C57BL/6J male and females (8-14 weeks old) sourced from Charles River (Cambridge, UK, RRID:IMSR_JAX:000664). Animals were housed in in temperature-controlled rooms (21°C) with a 12-hr light/dark cycle in groups of up to eight with sufficient enrichment, nesting material and water and food ad libitum. Sacrifice of animals was done so by a combination of either cervical dislocation or rising concentration of CO_2_ and then confirmation by exsanguination by severance of the femoral artery.

#### *In silico* analysis of colonic sensory neuron single-cell RNA sequencing dataset

The scRNA-seq database published by Hockley *et al*., 2019 (https://hockley.shinyapps.io/ColonicRNAseq/) was examined for the transcript expression of a variety of genes associated with the canonical IL6/OSM signalling pathway (Figures 1 and 2) Expression of a gene was defined as any transcription (log TPM) >0.

**Figure 1.**
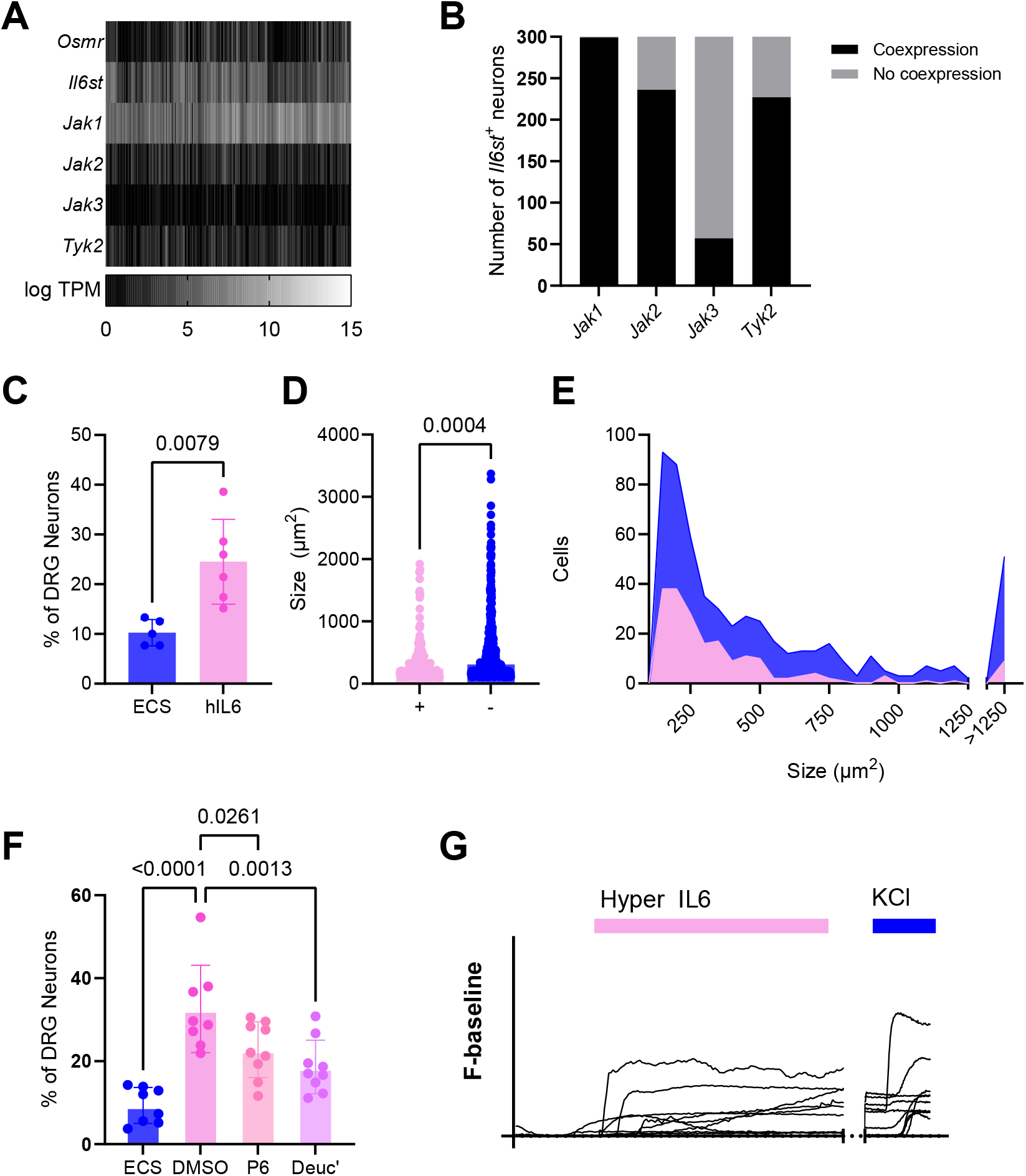
Hyper IL6 evokes [Ca^2+^]_i_ increase in DRG Neurons. (A) Heatmap showing the expression (logTPM) of *Osmr, IlCst, Jak1, Jak2, Jak3* and *TYK2* in murine colonic sensory neurons (data redrawn from Hockley et al. 2019). (B) Stacked bar chart depicting the proportion of *IlCst*^*+*^ neurons co-expressing *Jak1, Jak2, Jak3* and *Tyk2* (data redrawn from Hockley et al. 2019). (C) The proportion of DRG neurons responding to vehicle (ECS; extracellular solution) and 1 minute of 3nM Hyper IL6 stimulation (ECS (n = 5) vs. 3nM Hyper IL6 (n = 6): *p =* 0.0079, two-tailed Welch’s t-test; t = 3.879, df = 6.108). (D) Scatter dot plot comparison of cell soma sizes (μm^2^) between Hyper Il6 responders (+) and non-responders (-) (*p* =0.0004, two-tailed Mann-Whitney test; *U* = 45074). (E) Cell size distribution of responders and non-responders to Hyper Il6. (F) The proportion of DRG neurons responding to 3nM Hyper Il6 following pre-incubation with DMSO (vehicle for antagonists), 1μM P6 (Pyridone 6; Pan-JAK inhibitor) or 100nM Deuc’ (Deucravacitinib; TYK2 allosteric inhibitor) (*p* = 0.0261 and *p* = 0.0013, respectively, ordinary one-way ANOVA (main effect: *p* = <0.0001; F (3, 30) = 13.98) with Dunnett’s multiple comparison; *n* = 8, *N* = 8 and *n =* 9, *N* = 9). (G) Representative traces of Fluo-4 fluorescence during the application of 3nM Hyper Il6 and 50mM KCl from 10 neurons (break in the x-axis represents a 4-minute period of ECS wash).

**Figure 2.**
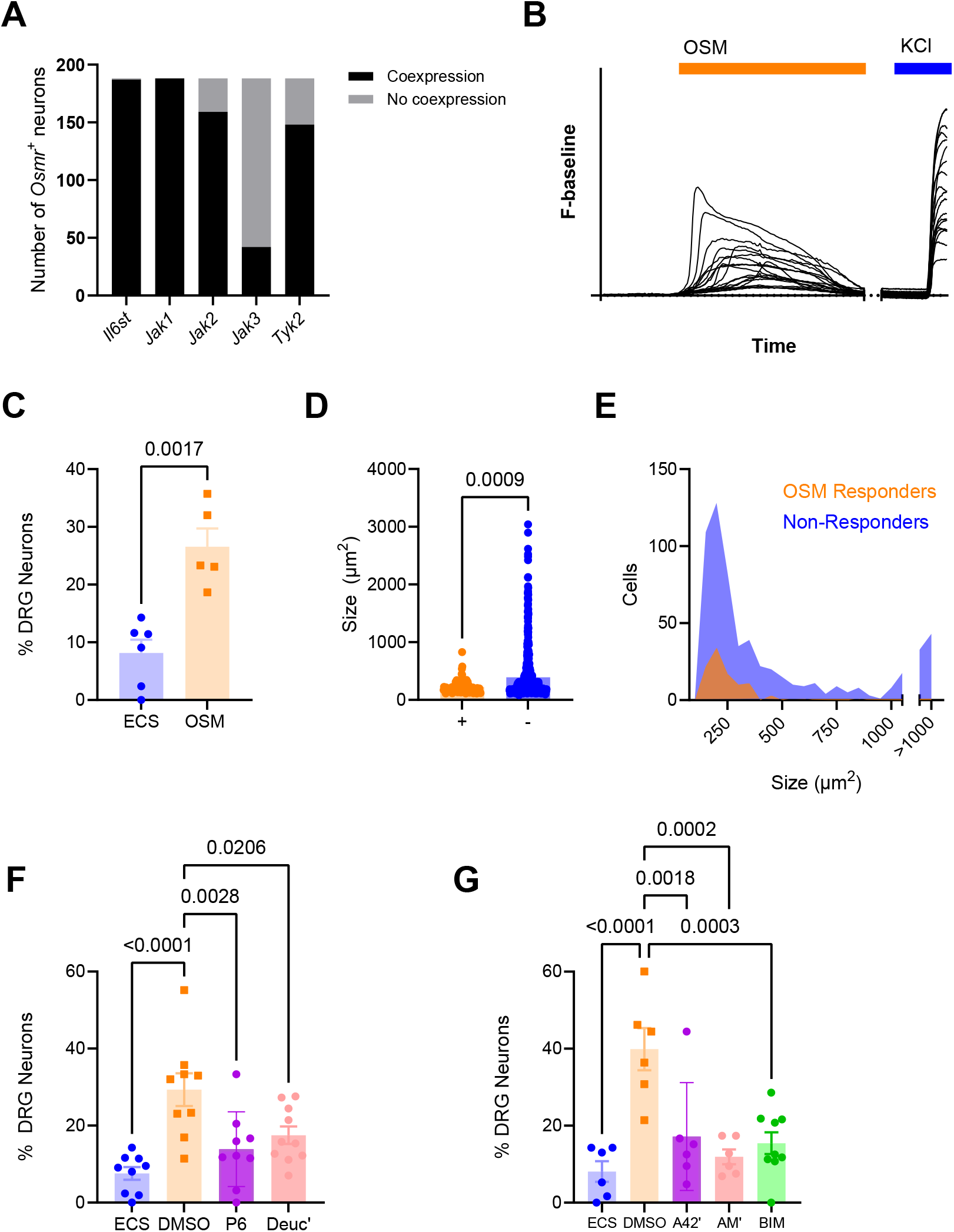
Oncostatin M (OSM) evokes [Ca^2+^]_i_ increase in DRG Neurons, mediated by TYK2. (A) Stacked bar chart depicting the proportion of *Osmr*^*+*^ neurons co-expressing *IlCst, Jak1, Jak2, Jak3* and *Tyk2* (data redrawn from Hockley et al. 2019). (B) Representative traces of Fluo-4 fluorescence during the application of 30nM OSM and 50mM KCl from 19 neurons (break in the x-axis represents a 4-minute period of ECS wash). (C) The proportion of DRG neurons responding to vehicle (ECS; extracellular solution) and 1 minute of 30nM OSM stimulation (ECS (n = 6) vs. 30nM OSM (n = 5): *p =* 0.0017, two-tailed Welch’s t-test; t = 4.709, df = 7.695). (D) Scatter dot plot comparison of cell soma sizes (μm^2^) between OSM responders (+) and non-responders (-) (*p* =0.0009, two-tailed Mann-Whitney test; *U* = 22247). (E) Cell size distribution of responders and non-responders to OSM. (F) The proportion of DRG neurons responding to 30nM OSM following pre-incubation with DMSO (vehicle for antagonists), 1μM P6 (Pyridone 6; Pan-JAK inhibitor) or 100nM Deuc’ (Deucravacitinib; TYK2 allosteric inhibitor) (*p* = 0.0028 and *p* = 0.0206, respectively, ordinary one-way ANOVA (main effect: *p* = 0.0002; F (3, 33) = 9.134) with Dunnett’s multiple comparison; *n* = 9, *N* = 9 and *n =* 10, *N* = 10 respectively). (G) The proportion of DRG neurons responding to 30nM OSM following pre-incubation with DMSO, 1μM A425619 (TRPV1 antagonist), 1μM AM0902 (TRPA1 antagonist), and 1μM BIM (Bisindolylmaleimide I, PKC inhibitor) (*p* = 0.0018, *p* = 0.0002, and *p* = 0.0003 respectively, ordinary one-way ANOVA (main effect: *p* = <0.0001; F (4, 28) = 9.382) with Dunnett’s multiple comparison; *n* = 6, *N* = 6 and *n =* 8, *N* = 8).

Heat maps demonstrating the relative quantity of gene transcripts for various genes, as well as co-expression profiles, were generated using GraphPad Prism software version 10.6.1 (892) using data derived from the above database.

#### Dorsal root ganglia sensory neuronal cell dissociation and culture

Dorsal root ganglia (DRG) from the levels which innervate the distal colon (T12-L5) were extracted from the murine spine and underwent axotomy before a process of enzymatic dissociation. DRG were incubated at 37°C with 5% CO_2_ in a media of 6-mg·ml^−1^ bovine serum albumin (BSA) in Leibovitz’s L-15 Medium and GlutaMAX™ Supplement with an additional 2.6% [v/v] NaHCO_3_ 1-mg·ml^−1^ which was then supplemented with initially type 1A collagenase for 15 mins, followed by 1-mg·ml^−1^ trypsin for 30 mins. DRG were then transferred to a L-15 media and GlutaMAX™ Supplement with added 10% (v/v) foetal bovine serum, 2.6% (v/v) NaHCO_3_, 1.5% (v/v) D-glucose where they underwent mechanical dissociation by repeated gentle titruation using a P1000 pipette with regular bore tip. Between each round of trituration, the DRG were pelleted via centrifugation (100xg for 1 minute) and the supernatant containing dissociated cells was collected. Following four rounds of trituration the supernatant underwent centrifugation for five minutes at 300xg, the supernatant removed and the pellet resuspended in the residual volume. This cell suspension was then seeded onto laminin-coated and poly-D-lysine–coated, 35mm dishes (MatTek, Ashland, MA) and incubated at 37°C in 5% CO_2_ for 1.5 hours to ensure cell adhesion to the dish before being flooded with 2ml of the D-glucose-supplemented media stated above for 16-24 hours prior to imaging.

##### Ca^2+^ imaging of cultured sensory neurons

The calcium indicator Fluo-4-AM was loaded into cells by incubation at 10 µM in extracellular solution (ECS; 140 mM NaCl, 4 mM KCl, 1 mM MgCl_2_, 2 mM CaCl_2_, 4 mM D-glucose, and 10 mM HEPES; pH 7.35-7.45) for 40 ± 5mins at room temperature and protected from light. Following incubation, cells were washed with ECS and then 200 µl ECS added before the dish is mounted onto the stage of an inverted microscope (Nikon Eclipse TE2000S). In experiments where an antagonist is used, following Fluo-4-AM incubation, 200 µl of the drug solution was applied to the cells and incubated for 10 mins at room temperature and shielded from light. Antagonist was then removed and 200 µl of ECS added. All drugs superfused onto the cells were diluted to the final concentration in 5 ml ECS and superfused at a rate of ∼0.5 mL/minute using a gravity-fed perfusion tip positioned adjacent to the field of view.

Recordings were taken with a CCD camera (Retiga Electro, Photometrics, Tucson, AZ) at 2.5 Hz with 100 ms exposure. Fluo-4-AM was excited by a 470 nm light source (Cairn Research, Kent, UK), and recordings taken at an emission wavelength of 520 nm and captured using open-source µManager software developed by Stuurman *et al*, (2010). Where multiple drugs were added to the same dish, a 4 minute period of ECS superfusion occurred between applications. At the end of each experiment, 50 mM KCl was applied to identify viable neurons and enable the normalisation of fluorescence.

Image analysis was performed using ImageJ. Neurons were designated as regions of interest by manual drawing around the perimeter of the cell body, with average pixel intensity per cell per frame measured.

Analysis consisted of the subtraction of background fluorescence from the dish (a defined cell-less region), and the measured region fluorescence (F) normalised to a 10 second (25 frames) period of ECS superfusion prior to drug application. Any cell which exhibited an increase in fluorescence of >5% relative to the maximum fluorescence (F_pos_) measured during 50mM KCl application was taken forward for further analysis. To determine a cell as a positive responder to any particular drug/vehicle, a stable baseline and an increase in fluorescence of >0.1 F/F_pos_ was required.

#### Statistics

Normality of data was tested using Shapiro-Wilk normality test (alpha = 0.05) and, when necessary, the appropriate non-parametric test was used. GraphPad Prism software version 10.6.1 (GraphPad Inc, La Jolla, CA) was used to undertake all statistical analyses, with specific tests used detailed in the corresponding figure legend.

## Results

### *Osmr, IlCst*, and the Janus Kinases are expressed in colonic sensory neurons

By utilising the single-cell transcriptomic dataset of colonic sensory neurons generated by Hockley *et al*. (2019) it is possible to examine the expression of genes encoding OSMR (*Osmr*), gp130 (*Il6st*), and the Janus Kinases (*Jak1, Jak2, Jak3*, and *Tyk2*). Broad expression profiles are apparent for *Il6st* and *Jak1*, with more restricted but nonetheless substantial expression of *Osmr, Jak2*, and *Tyk2* (Figure 1A.) Whilst OSM requires the presence of gp130 and OSMR on the cell membrane, Hyper Il6 signalling only needs to bind to gp130 to induce autophosphorylation of a heteromeric dimer of Janus Kinases. When the *Il6st*^*+*^ population is examined for co-expression with any of these JAKs, it can be see that although *Jak1* is expressed in 302 out of the 303 *Il6st*^*+*^ neurons, both *Jak2* and *Tyk2* are also expressed in 87.8% and 81.8% of *Il6st*^*+*^ neurons (Figure 1B.) suggesting that a large proportion of colonic sensory neurons possess the necessary machinery to transduce a Hyper IL6 signal intracellularly.

### Hyper Il6 elicits a rise in intracellular calcium in DRG neurons

Using the cell bodies of primary neurons extracted from the DRG of mice, an attempt was then made to establish a functional role for gp130 in the depolarisation of nociceptive populations by observing for an increase in cytosolic Ca^2+^ ([Ca^2+^]) in response to hyper Il6 stimulation.

Of total neurons imaged, 24.51 ± 8.53% were found to respond to 1 minute superfusion of 3nM hyper Il6 (93 neurons from a total of 370 from six mice; Figure 1C.). When the size of hyper Il6 responders is compared with that of non-responsive neurons, it is apparent that the size of hyper Il6 responders is significantly smaller and predominantly in the 110-800 μm^2^ soma size range, indicative of nociceptors (grouped data of neurons from 12 dishes from six mice; Figures 1D. and E.). These findings suggest that gp130 activation can induce an increase in cytosolic Ca^2+^ and may be occurring in a nociceptive subpopulation.

#### Hyper Il6-induced calcium mobilisation involves TYK2 activation

To further examine this signalling, neuronal cultures were pre-incubated with either vehicle (DMSO), a pan-JAK antagonist (Pyridone 6; 1μM) or a TYK2-specific allosteric inhibitor (Deucravacitinib; 100nM) prior to hyper Il6 stimulation. As apparent in Figure 1F. disruption of all JAKs does reduce the proportion of responsive neurons to hyper Il6 (*p* = 0.0261, 32.6 ± 10.52% vs. 22.81 ± 6.72% respectively) however, inhibition of TYK2 alone appears sufficient to induce a similarly significant reduction in responsive neurons (*p* = 0.0013, 32.60 ± 10.52% vs. 18.63 ± 6.45% respectively).

Whilst these data provide an interesting insight into gp130 signalling, for other Il6-family cytokines another membrane bound receptor is required for intracellular signalling. Hyper Il6 is designed to mimic the action of Il6 and the Il6 soluble receptor forming a complex with gp130, a mechanism by which IL6 can signal in cells that don’t express the membrane bound Il6 receptor -including neurons. Due to the somewhat artificial nature of this Il6-Il6 receptor fusion protein, it is also important to examine this signalling mechanism using a naturally endogenous Il6 family cytokine such as OSM.

#### OSM also increases cytosolic Ca^2+^ in DRG neurons

As demonstrated in Figure 2A., *Il6st* is expressed in all but one of the 238 *Osmr*^+^ cells, and similarly to within the *Il6st*^*+*^ population, *Jak2* and *Tyk2* are expressed in high proportions (91.6% and 84.9% respectively) alongside *Jak1* which is expressed in 100% of the *Osmr*^*+*^ population. It would therefore be expected that perfusion of OSM would also elicit an increase in cytosolic Ca^2+^. 1 minute of 30nM OSM perfusion was shown to activate 26.55 ± 7.05% compared to 8.14 ± 5.68% which did so in response to ECS alone (Figure 2C.). Concurrent with the idea of that small diameter neurons are the predominant group of OSM-responders, measurement of cell soma area shows the majority or responders lie within the 150 - 500μm^2^ range, indicative of a nociceptor population (Figures 2D. and E.).

As was done with hyper Il6, cells were pre-incubated with either vehicle (DMSO), Pyridone 6 (P6; 1μM) Deucravacitinib (Deuc’;100nM) before application of 30nM OSM. The proportion of responders to 30nM OSM + DMSO was found to be similar that found in the previous group with an average of 29.33 ± 12.72%, which was then reduced when all JAKs were inhibited (13.89 ± 9.70%) or when only TYK2 was blocked (17.49 ± 7.23%; Figure 2F.). This further supports the hypothesis that OSM-induced calcium mobilisation specifically involves TYK2 activation.

The mechanism by which OSM would elicit a change in [Ca^2+^] is unknown, however, considering the sensitisation observed of TRPV1 by other receptor tyrosine kinases -most notably nerve growth factor (NGF)-this may be a route by which OSM operates (Jin *et al*., 2004; Stein *et al*., 2006). To examine this, prior to OSM superfusion, cells were incubated with the potent and highly specific TRPV1 antagonist A425619 (1μM), followed by OSM superfusion as previously described (El Kouhen *et al*., 2005). As observable in Figure 2G., the proportion of OSM-responsive neurons was significantly reduced in the presence of a TRPV1 antagonist (39.86 ± 13.42% vs 17.17 ± 14.02%, p< 0.01).

This protocol was done in the presence of an antagonist against TRPA1, Similarly, TRPA1 - which is also highly expressed in murine colonic sensory neurons (Hockley *et al*., 2019)-antagonism produced a significant reduction in the proportion of OSM-responsive neurons (39.86 ± 13.42% versus 11.91 ± 4.71, p<0.01; Figure 2G.). This correlates with that reported by Malsch *et al*. (2014) where TRPA1-mediated calcium transients were seen to be attenuated in *gp130*^*-/-*^ Na_v_1.8-expressing small diameter DRG neurons, as well as a marked decrease in *Trpa1* expression in both *gp130*^*-/-*^, and *gp130*^*ff/ff*^ cells incubated with antagonists against all janus kinases and STAT transcription factors. Conversely, a similar decrease in the proportion of capsaicin-responsive neurons was not observed, although the effect on calcium-transient magnitude was not reported.

This finding raises the further question of how OSM signalling may result in TRPV1 and TRPA1-mediated calcium entry. Previous studies on these channels have highlighted the phosphorylation of Protein Kinase C (PKC) as a factor in the sensitisation of both TRPV1 and TRPA1 in sensory neurons and classically activated downstream of the inflammatory mediator bradykinin (Vellani *et al*., 2001 Wang *et al*., 2008; Brackley *et al*., 2017). PKC is also found to be activated downstream of OSM-JAK-PI3K signalling and so could be a player in the observed increase in cytosolic calcium in response to OSM exposure (Smyth, Kerr and Richards, 2006). When 30nM OSM was applied in the presence of a pan-PKC inhibitor bisindolylmaleimide I (BIM; 1μM), there was a significant decrease in the proportion of OSM-responsive DRG neurons (39.86 ± 13.42% versus 15.44 ± 8.52%, p< 0.001; Figure 2G.).

## Discussion

The dearth of effective analgesics for patients with IBD continues to reduce quality of life and drive sufferers into the use of clinically dubious opioid options (Kurlander and Drossman, 2014). Understanding the way in which factors ubiquitous in diseased tissue, such as pro-inflammatory cytokines, contribute to nociception in the gut is vital for advancing the development of effective alternative pain treatments. This study intended to determine a role for TYK2 in IL-6 family cytokine signalling in nociceptive sensory neurons. By examining changes in cytosolic calcium in murine DRG neurons we show that increased [Ca^2+^]_i_ elicited by hyper IL6 and OSM can be attenuated by pharmacological inhibition of all JAKs but importantly, by allosteric inhibition of TYK2 alone.

We observed an increased proportion of sensory neurons showing [Ca^2+^]_i_ in response to hyper IL6 and OSM superfusion, with inhibition of downstream JAKs and non-selective ion channels TRPV1 and TRPA1 (for OSM). This provides useful insight into the mechanism by which this increase in intracellular calcium was occurring however, this does not necessarily conclude that hyper IL-6 and OSM are acting directly on these responsive cells. As primary DRG cultures contain other non-neuronal cells such as glia and satellite cells, it is possible that they may be acting upon these, with subsequent secondary mediators eliciting calcium transients in neurons. Expression of OSMR was not previously observed in satellite cells in L1-L5 murine DRG, which would suggest a similar transcriptome in resident immune cells within the other spinal levels examined in this study (Tamura *et al*., 2003). There is supportive evidence for OSM acting directly on the activated neurons as the size of *Osmr*^*+*^ neurons measured by Tamura *et al*. (2003) and Morikawa *et al*. (2004) showed similarity in the small soma size neurons as we observed here. The latter study by Morikawa *et al*. (2004) also demonstrated that *Osmr*^*-/-*^ mice showed less nocifensive behaviour to capsaicin intraplanar injection compared to wild-type, suggesting a role for OSM signalling in the function of TRPV1, with a reduction in the proportion of TRPV1^+^ neurons in the DRG. Equally, mechanical hypersensitivity and TRPA1 expression has been observed to be reduced following the deletion of *gp130*, corroborating our implication of TRPA1 in hyper Il-6 and OSM [Ca^2+^]_i_ flux (Malsch *et al*., 2014).

A recent study by Li *et al*. (2026) also substantiated a nociceptive function for OSM in small diameter neurons in rats, however they observed little co-sensitivity between OSM responsive and capsaicin responsive neurons and a lack of thermal hypersensitivity following intrathecal OSM injection. This disparity is likely a result of species differences.

The question as to how OSM acts in conjunction with TRPV1, TRPA1 and potentially any alternative mechanosensitive ion channels is one which requires further investigation, however the attenuative effect observed when PKC was inhibited suggests this may play a role. TRPV1 has previously been shown to be sensitised via PI_3_K and PKC-δ, as well as OSM increasing pERK and PKA-pRII levels, leading to hyperalgesic priming (Andrarsch *et al*., 2009; Garza Carbajal *et al*., 2020). The intricate interplay between various signalling molecules could provide many possible targets for pharmacological intervention, although ensuring there are limited immunological side-effects when inhibiting pro-inflammatory pathways is crucial for clinical applications.

We have used calcium imaging to demonstrate a role for TYK2 in OSM signalling in the activation of sensory neurons via TRPV1 and TRPA1. This highlights a novel target for targeting OSM-mediated nociception which could have implications in the treatment of inflammatory pain, such as that in IBD.

This understanding will facilitate further exploration of the mechanisms by which nociception in IBD occurs and highlight options for *in vivo* animal studies to examine the role OSM signalling plays in the context of the myriad of inflammatory pathways occurring in the diseased colon.

